# Functional Control of Network Dynamical Systems: An Information Theoretic Approach

**DOI:** 10.1101/2024.06.17.599263

**Authors:** Moirangthem Sailash Singh, Ramkrishna Pasumarthy, Umesh Vaidya, Steffen Leonhardt

**Affiliations:** Electrical Department, Indian Institute of Technology, Madras, Chennai, 600036, Tamil Nadu, India; Mechanical Department, Clemson University, 105 Sikes Hall, Clemson, 29634, South Carolina, USA; Helmholtz Institute of Biomedical Engineering, RWTH Aachen University, Pauwelsstr, Aachen, 52074, Germany

## Abstract

In neurological networks, the emergence of various causal interactions and information flows among nodes is governed by the structural connectivity in conjunction with the node dynamics. The information flow describes the direction and the magnitude of an excitatory neuron’s influence to the neighbouring neurons. However, the intricate relationship between network dynamics and information flows is not well understood. Here, we address this challenge by first identifying a generic mechanism that defines the evolution of various information routing patterns in response to modifications in the underlying network dynamics. Moreover, with emerging techniques in brain stimulation, designing optimal stimulation directed towards a target region with an acceptable magnitude remains an ongoing and significant challenge. In this work, we also introduce techniques for computing optimal inputs that follow a desired stimulation routing path towards the target brain region. This optimization problem can be efficiently resolved using non-linear programming tools and permits the simultaneous assignment of multiple desired patterns at different instances. We establish the algebraic and graph-theoretic conditions necessary to ensure the feasibility and stability of information routing patterns (IRPs). We illustrate the routing mechanisms and control methods for attaining desired patterns in biological oscillatory dynamics.

**Author Summary:** A complex network is described by collection of subsystems or nodes, often exchanging information among themselves via fixed interconnection pattern or structure of the network. This combination of nodes, interconnection structure and the information exchange enables the overall network system to function. These information exchange patterns change over time and switch patterns whenever a node or set of nodes are subject to external perturbations or stimulations. In many cases one would want to drive the system to desired information patterns, resulting in desired network system behaviour, by appropriately designing the perturbating signals. We present mathematical framework to design perturbation signals that drive the system to the desired behaviour. We demonstrate the applicability of our framework in the context of brain stimulation and in modifying causal interactions in gene regulatory networks.

## 1 Introduction

Recent advancements in brain stimulation techniques offer new possibilities for diagnosing [1], monitoring [2], and treating neurological and psychological disorders [3]. For instance, non-invasive techniques such as transcranial magnetic stimulation (TMS) and transcranial direct current stimulation (tDCS) are utilized to treat conditions like epilepsy [4], attention deficit hyperactivity disorder (ADHD) [5], schizophrenia [6], and tinnitus [7]. In contrast, invasive methods such as deep brain stimulation (DBS) are employed to treat dystonia, essential tremor, medically resistant epilepsy, Parkinson’s disease, and medication-resistant obsessive-compulsive disorder (OCD) [8].

Despite its broad applications, various key challenges remain on the optimal stimulation parameters, identification of target areas that maximize clinical utility [9] and minimizing stimulation-induced side effects [10]. To determine the optimal parameters and target areas, methods have been developed to use the information from imaging techniques (CT scan, fMRI, fast optial imaging) and recording devices (PET, MEG, EEG etc) to monitor the effects of brain stimulation. Moreover, concepts of average controllability and modal controllability from network control theory, have been used to predict whether the effects of stimulation remain focal or spread globally [11, 12]. However, conventional brain stimulation techniques frequently cause undesirable side effects by inadvertently stimulating adjacent brain structures, thus limiting its effectiveness. To mitigate these side effects, one approach is to design electrodes that enable directional stimulation, thereby avoiding neighboring structures [13, 14]. An alternative approach is to design control strategies that facilitate precise directional stimulation at desired instances.

To this end, we propose an information-theoretic based functional connectivity, defined by the directional flow of information or causal inferences among neurons across the brain network. The functional connectivity holds fundamental significance, as biological network systems depend on the dynamic communication and exchange of information among cells and their associated subsystems. For instance, in gene-regulatory networks, information flow quantifies a cell’s capacity to regulate the protein concentrations of other cells [15], taking into account the inherent randomness of individual molecular events. In neurological networks [16], information flows across synapses through the coordinated activity of multiple neural populations. Dendrites carry information towards the cell body, while axons transmit it away from the cell body. The information flow is defined as the influence of the excitation level of an excitatory neuron on the neighbouring neurons. Therefore, understanding the information flows among neurons or cells offer insights into the patterns of information routing associated with various brain activities, potentially introducing a new dimension in the treatment of neurological or psychiatric disorders. The significance of information transfer in brain stimulation techniques lies in its ability to describe how the directional and quantitative influence of the stimulation propagates across the entire network.

Neurons in neurological networks, proteins in gene regulatory networks, and cells in various biological networks are interconnected through a network of oscillators. These nodes exhibit oscillatory and synchronous dynamics, often incorporating a stochastic component [17–21]. Therefore, understanding the dynamics of how information flows among the nodes in these complex networks offers a fundamental challenge because of their inherent complexity and non-linear dynamics. The results in [22, 23] demonstrate that oscillatory dynamics facilitate the transmission of information in biological and oscillatory networks. These results lead to a fundamental question: How do the patterns of information routing across the network vary based on the intrinsic dynamics of nodes, and external driving signals or simulations? The results in [24] provide insights into how information flow is influenced by changes in network topology and noise. An important result in [22, 25] provides a fundamental mechanism illustrating how fluctuations in the phase differences of multiple oscillators give rise to vaarious IRPs. This mechanism also elucidates how these IRPs can adaptively transition between multiple stable states under the influence of external inputs. The phenomenon is particularly relevant in the context of neurological networks, where the information flows or the causal interactions among different brain areas are reconfigured to achieve various brain functions such as vision, memory, or motor preparations [26]. Importantly, these transitions between multiple stable states take place due to the combined influence of structured “brain noise” and the bias imposed by sensory and cognitive driving, even when the underlying structural (anatomic) connectivity remains constant.

The primary focus of this study is on examining how the direction and magnitude of causal interactions among nodes change under external stimulations while keeping the structural connectivity fixed. We employ network control theory and information theory to design optimal control inputs or stimulations, enabling flexible “on-demand” selection of functional patterns.In the context of brain stimulation, the control policy aims to find the optimal inputs that achieve the desired functional pattern, defined by the intended direction and magnitude of excitation levels among connected neurons across the brain network when different regions are stimulated. The study also provides valuable insights into the optimal energy required for the transition from one information routing pattern to another. Towards this goal, we provide the conditions on the network dynamics and coupling strengths among the oscillators under which one can optimally reroute information routing patterns to a desired pattern. Related literature on the control of brain networks includes using control techniques to find the optimal trajectories to steer from an initial state (baseline condition) to target states (high activity in sensorimotor systems) in finite time and with limited energy [27, 28]. These studies focus on the control of state trajectories, and there remains a limited body of research dedicated to the control of information flows or routing patterns in complex dynamical networks. One of the few works in the control of functional patterns (described by simple statistical dependencies or correlations among oscillators) is studied in [23, 29]. Other studies on the control of brain network [30–35] focus on examining the ease of accessibility for specific states within a dynamic regime, identifying the regions that require perturbation to enable access to these states, and quantifying the energy required to attain them.

We first demonstrate how the dynamic state of oscillatory networks, and the presence of noise collectively give rise to a distinct communication pattern. This pattern is quantified using an information-theoretic measure defined by Kleeman’s Information transfer [36, 37]. Our selection of this information transfer definition is based on its rigorous derivation, as presented in [36, 37], which enables the determination of instantaneous information flows. Furthermore, the formulation has gained widespread adoption in numerous applications such as in financial markets [38, 39], in studying climate science [40], and for detecting causality in time series data [41, 42]. We show that information routing patterns or functional patterns depend on the underlying synchronized or equilibrium states and switching between multiple stable states generates multiple information routing patterns. The result enables the study of network control theory for non-linear systems around the stable equilibrium states. We develop a mathematical framework to determine the optimal energy levels needed for external driving signals to facilitate transitions among the information routing patterns. We demonstrate how interventions in the form of noise and external inputs at specific nodes within the network can alter the flow of information between other nodes. The results provide generic insights into the mechanisms underlying information rerouting in complex networked systems. We also illustrate how our framework can be expanded to redirect information routing patterns towards desired configurations, both in finite and stationary time horizons.

## 2 Results

### 2.1 Information Routings in Coupled Oscillators

The study of brain networks, gene regulatory networks, and various other biological networks can be conducted within the framework of complex networks with oscillatory dynamics. Therefore, to understand how information transfer patterns change dynamically under the influence of interactions between nodes, we consider a network of *n* coupled oscillators evolving according to

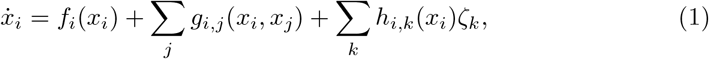

with *x*_*i*_ ∈ ℝ^*n*^, *f*_*i*_ smooth vector fields, *g*_*ij*_ coupling functions, and *h*_*i,k*_ denotes the impact of the random processes *ζ*_*k*_ with zero mean. We focus on weakly coupled oscillators where the separation between the phase and amplitude dynamics is possible [43]. Moreover, when the couplings are weak, we can reduce the system of nonlinear equations to a set of equations on a torus using invariant manifold theory [43]. Under this assumption, the phase-amplitude interactions among the individual nodes are negligible and the phase-amplitude dynamics are decoupled. Thus the network oscillatory dynamics can be expressed in terms of nodal phases only [44]. The stochastic oscillatory dynamics in equation (1) can thus be reduced to the averaged phase stochastic dynamics as

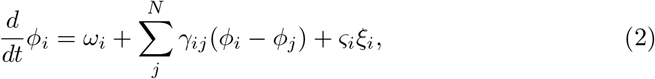

where *ω*_*i*_ denotes the intrinsic frequencies of node *i, γ*_*i,j*_ denotes the coupling function that depends on the phase differences only, and *ς*_*i*_ is a Gaussian noise process, *w*_*i*_ with zero mean and 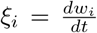. Supporting information B provides a comprehensive derivation of equation (2) from equation (1).

We decompose the phase dynamics into a deterministic reference part, 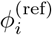, and a fluctuating component, 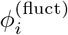 and focus on the phase-locked configuration with constant phase offsets, 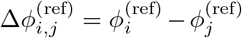. The fundamental idea is that the information routing patterns are determined by the fluctuating state variables around the stable phase locking states. Note that, in oscillatory networks, there may exist multiple reference deterministic solutions, giving rise to different IRPs among the nodes. The fluctuating component 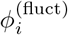 is estimated using a small noise expansion giving a first-order approximation for the evolution of the fluctuating component of the phase dynamics [22] (Methods and Supporting information B). Although the approximation around the stable phase-locking states might not precisely represent the information transfers among the oscillatory nodes, it offers a valuable benefit by providing an analytical approximation for the evolution of probability distributions defined by *ρ*_*j*|*i*_, *ρ*_*;j*_ in equation (23) (Methods). The analytic model derived from the linearized model provides a close approximation to the underlying functional connectivity of oscillatory networks [22]. Supporting information B provides its relevance in the context of weakly connected oscillatory networks, such as those found in neurological systems. Further, we assume that the noise levels remain sufficiently low to prevent transitions between multiple stable states. However, this assumption can be relaxed, and we can compute and control the information transfer function for each switching instant. In this case, the network becomes temporal (time-varying). Our assumption for the white noise model has the advantage of being tractable, at least in providing some simple differential equations for the evolution of probability distributions. Using the formulation in equation (23) (Methods) together with the first-order approximation, information transfer from state *x*_*j*_ to *x*_*i*_ at time *t* is derived as (Supporting information A)

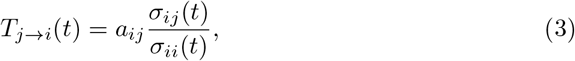

Where 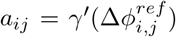 and *σ*_*ij*_ denotes the (*i, j*) component of the state covariance matrix at time *t*, ∑(*t*). Using equation (3), the IRP for a network of *n* oscillators at time *t*, denoted as IRP^*t*^ can be written as IRP^*t*^(*i, j*) = *T*_*j*→*i*_(*t*). Thus, given a network of *n* coupled oscillators with coupling function *γ*_*ij*_ and initial covariance ∑(0), IRP^*t*^ is an *n × n* matrix with IRP^*t*^(*i, j*) = *T*_*j*→*i*_(*t*), *i, j* ∈ {1, *… n*}. The IRP satisfies the relation (Theorem 1, Supporting information C)

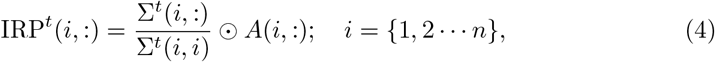

where ⊙ denotes the Hadamard or the element-wise matrix multiplication, ∑(*t*) is denoted by ∑^*t*^. The derivations in equations (23) and (3) are detailed in Supporting information A. The importance of concentrating solely on phase dynamics becomes apparent when considering that synchronization and phase-locking phenomena inherently foster interactions among the oscillators, acting as an innate catalyst for initiating the communication process. Similar investigations into phase dynamics are leveraged across a wide range of oscillatory networks to determine diverse functional relationships among the oscillators.

A model of a gene regulatory network consisting of two symmetrically coupled Goodwin oscillators (Supporting information D) is shown in Figure 1. Figures 1b-1e depict the IRP in the absence of any external input or influence. The deterministic reference states are determined by the zeros of the antisymmetric coupling 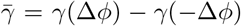 with negative slope as shown in Figure 2. The presence of an external signal and noise affecting one of the transcribed mRNAs induces fluctuations around one of these two stable states. This, in turn, results in a shift in the phase difference and eventually impacts the couplings between the two oscillators. Sufficiently strong external signals have the potential to induce transitions between the two stable states, facilitating the interchange in the direction of information flows between the two oscillators as shown in Figures 1f-1i, without even altering the structural properties of the network. Despite the symmetry of the oscillatory network, the asymmetrical information flows depicted in Figures 1e and 1i arise from the uneven coupling strengths in the phase dynamics. That is, for the two oscillators in Figure 1a, the evolution of the fluctuating component of the phase dynamics can be written from equation (2) as

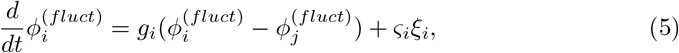

where the coupling strengths 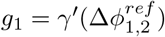 and 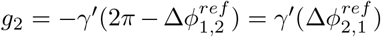. In the reference state 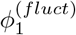, oscillator 1 (green) receives inputs from oscillator 2 proportional to 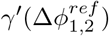, while oscillator 2 receives inputs from oscillator 1 propor-tional to 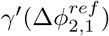. Since 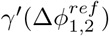 significantly deviates from the phase-locked state of oscillator 1, inputs to oscillator 2 exert a strong influence on oscillator 1 to restore the phase-locking state. The information transfer is thus dominant in the direction from oscillator 2 to 1 (*T*_*b*→*g*_, Figure 1e).

**Fig 1.**
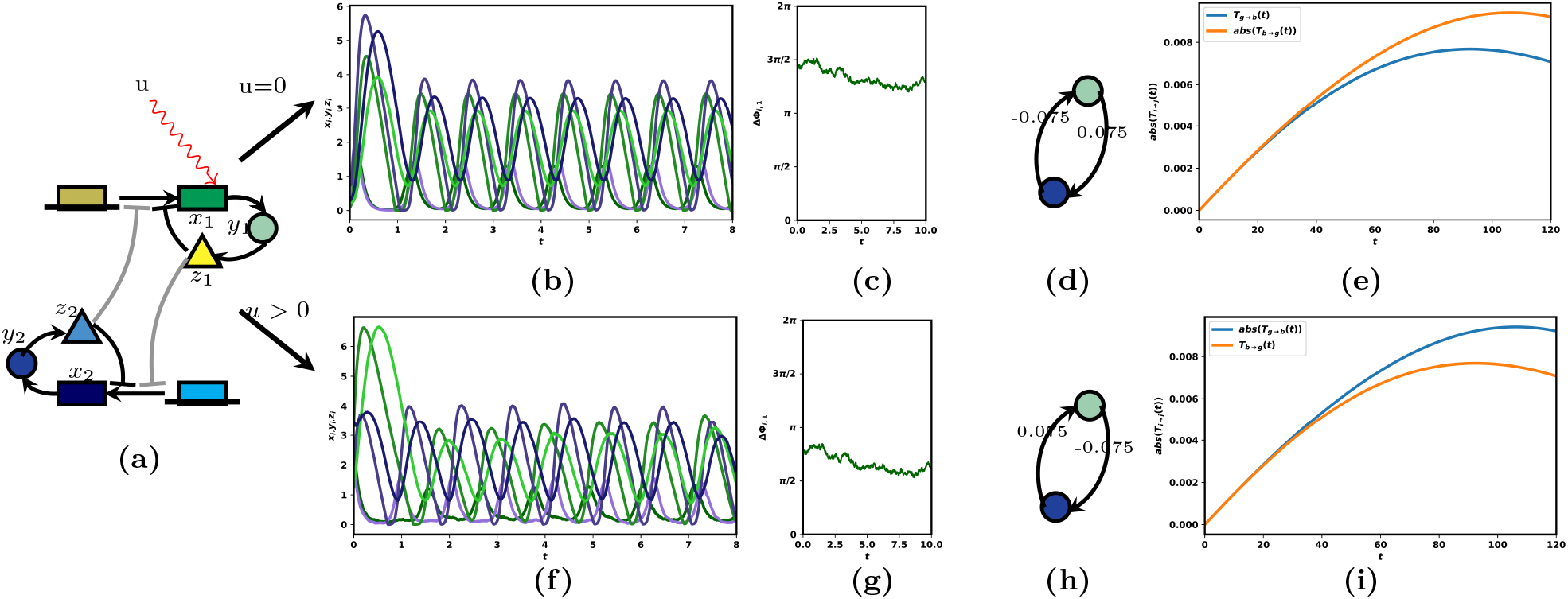
Impact of external signal on information routing patterns: (a) Simple model of a gene-regulatory network of two coupled Goodwin oscillators (yellow and blue). Coupling strengths are gray coded (darker color indicates stronger coupling), sharp arrows indicate activating, and blunt arrows inhibiting influences (b) oscillatory dynamics of the concentration levels of *x*_*i*_(mRNA), *y*_*i*_ (enzyme), and *z*_*i*_ (protein), *i* ∈ {1, 2}. (c) Fluctuation in the phase difference around a stable reference (d) Phase reduced network model around the stable phase locking state (e) Evolution of the information routings between the two oscillators (f) Same as in (b) but with influence from an external signal, *u* and (g) the resulting shift in the stable reference state (h) same as in (d) but due to the influence of *u* (i) change in the routing patterns due the influence of *u*.

**Fig 2.**
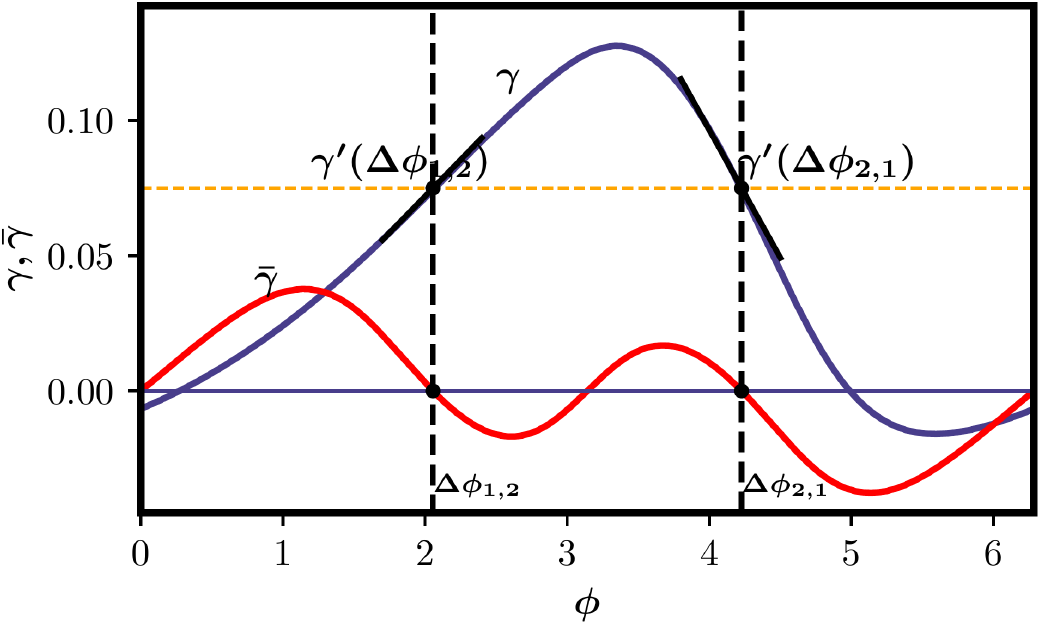
Phase Coupling function, *γ*(Δ*ϕ*) (blue) between the two oscillators and its antisymmetric coupling (red), 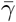 for the Goodwin Oscillator.

Note that the coupling strength from oscillator 2 to 1 in figure 1d is negative. This results in negative information transfer from oscillator 2 to oscillator 1, indicating that the influence from oscillator 2 to oscillator 1 is weakening, although it remains stronger in magnitude than the information flow from oscillator 1 to oscillator 2. With sufficiently strong input, the stable phase locking state is shifted as shown in figure 1g, resulting in the reverse directional information flow or pattern as shown in figure 1i. In the next section, we propose frameworks for steering information routing patterns toward a desired configuration in both finite time horizon and infinite time horizon scenarios.

### 2.2 Functional Control to achieve the desired IRP

The results in Figures 1 and 2 illustrate the significance of the coupling function’s behavior around the stable phase locking states in shaping the information routing patterns. The results in [24] elaborate on how multiplicative noise, additive noise, and local noise impact information transfer. This phenomenon wherein the functional properties of network systems are shaped by the coordinated behavior among their oscillatory components is present in diverse natural and technological network systems. For example, unique phase-locking patterns influence the coordinated movement of orbiting particle systems [45], facilitate successful mating in populations of fireflies [46], control active power flow in electrical grids [47], forecast global climate change phenomena [48], and support various cognitive functions in the brain [49, 50]. Despite its practical importance, the exploration and development of strategies to actively enforce specific information routing patterns have been relatively limited. In this section, we develop mathematical frameworks for designing optimal control inputs that reroute the IRPs to desired patterns. Figure 3 shows our framework and an example of control of information transfers in a network of 15 nodes at some finite instant *t* = 100.

**Fig 3.**
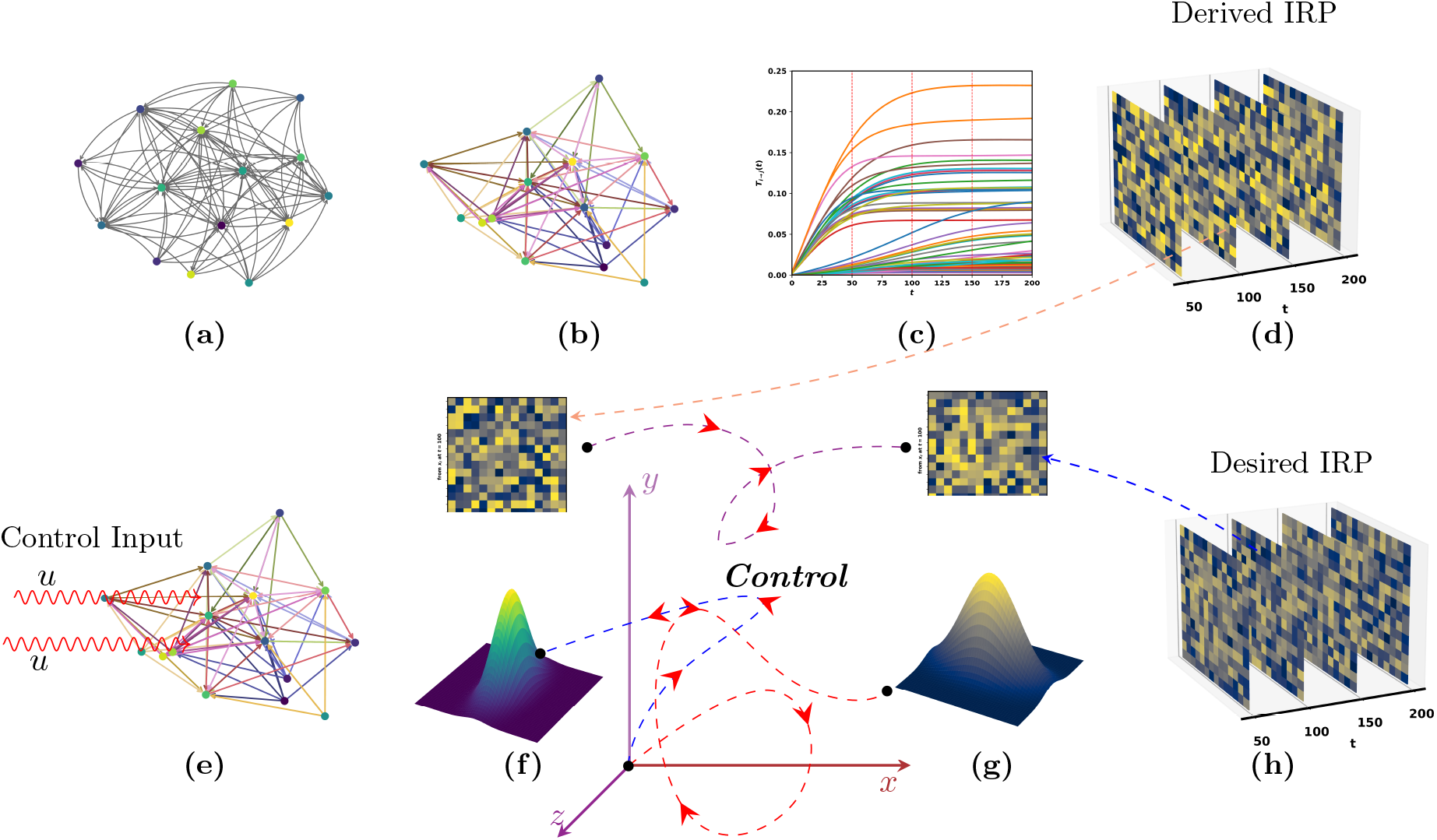
Network Control to achieve a desired information routing pattern from an undesired one: The figure shows a network of 15 nodes, with the initial network dynamics following a Gaussian distribution. (a) The structural network with oscillatory dynamics. (b) Phase-reduced network model for the oscillatory network in Fig. (a). (c) Evolution of the information transfers among the nodes derived using equations (23) and (3). (d) The derived IRPs using equation (4) at various time instances. (e) Introduction of control inputs on the phase-reduced network model to achieve a desired IRP at *t* = 100. (f) The probability distributions of the nodes and the derived IRP from Fig. (d) at *t* = 100. (g) Using the control inputs, the distribution corresponding to the desired IRP in Fig. (h) at *t* = 100 is achieved.

The concept of regulating IRPs differs across various applications, depending on the system objectives or performance. For example, in the context of brain stimulation techniques, the control inputs correspond to external stimulations applied to different brain regions, inducing additional fluctuations in the phase differences between these two neurons. Utilizing these control inputs, it becomes feasible to generate personalized control profiles for individuals. These profiles have the potential to improve our understanding of psychological and clinical impairments in patients, offering a more biologically and mechanistically informed perspective [51, 52]. The temporal changes in the neuronal response or the synaptic coupling strength during certain activities or learning processes [53] can lead to specific IRPs within a specified time frame. This represents an example of finite horizon control of IRP, where the learning processes and activities serve as the control input. In gene regulatory and cell-signaling networks [15, 54, 55], a specific IRP might be favored in an infinite time horizon through the process of evolution, achieved by influencing various states. In engineered non-oscillatory networks like a wireless networked control system, IRP defines the channel capacity of the communication channel, and controlling the IRP aligns with the control of the Signal-to-Interference-plus-Noise Ratio (SINR) in finite time. This expansion of scope highlights the versatility and applicability of the IRP concept, demonstrating its relevance in diverse domains beyond oscillatory networks. Considering these diverse applications, we define our IRP control problem in both the finite and infinite horizons.

The main objective of this section is to enforce a desired IRP within both finite and infinite time frames, as depicted in Figure 3. An approach to achieve this objective is to regulate the fluctuations around the stable phase locking states using state feedback control inputs. This approach is logical as it aligns with the fundamental concept of information routing patterns: *different IRPs arise due to the fluctuations in the phase differences among the oscillators [22]*.

With the control inputs *u*(*t*), the first-order approximation (Methods and Supporting information B) of the fluctuating component in equation (2) can be rewritten as 9

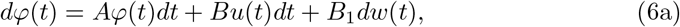

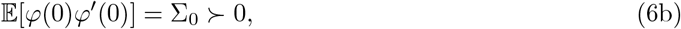

where *B* ∈ ℝ^*n×p*^ is the input matrix that specifies which of the oscillators are affected or influenced by the control inputs, and ∑_0_ is the initial state covariance matrix. The transpose of the state matrix, *A*′ ∈ ℝ^*n×n*^ describes the weighted adjacency matrix. The corresponding directed network is denoted by *𝒢* (*𝒱, ℰ*_*𝒜*_), with nodes *𝒱* = {1, 2, …*n*}, given by the *n* states, *ℰ*_*𝒜*_ = {(*i, j*)|*i, j* ∈ *𝒱*} is the edge set. We use the notation *U* to represent the collection of control functions with finite energy and define it as the set of permissible control inputs. We also use *X*′ to denote the transpose of a vector or a matrix *X*.

#### 2.2.1 Finite Horizon Minimum Energy Control of IRP

The objective is to achieve the desired IRP in a finite time horizon, *T*. The selection of the desired pattern at *T* is limited by the constraint on the positive definiteness of the state covariance matrix at *T*. Corollary 1.1 (Supporting information C) provides the admissible conditions for a given desired routing pattern. Given an admissible desired pattern IRP_*d*_, we can formulate the information routing control problem as

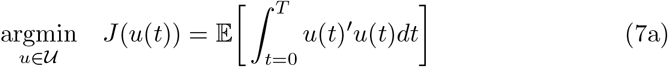

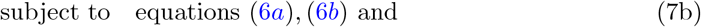

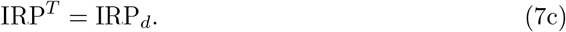

We show in Theorem 2, Supporting information C that the Problem (7) can be solved through suitable feedback control inputs if and only the network is controllable [56]. And given a controllable network [30], the control input solving Problem (7) is of the form *u*(*t*) = *K*(*t*)*ϕ*(*t*) (Proposition 1, Supporting information C). The optimal control strategy *u*^*^(*t*) solving Problem (7) is then of the form

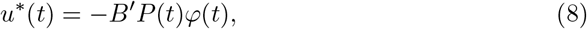

where *P* (*t*) is a differentiable matrix function taking values in the set of *n×n* symmetric matrices and satisfies both the following coupled Riccati equations:

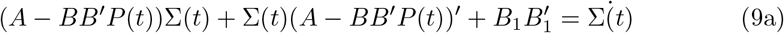

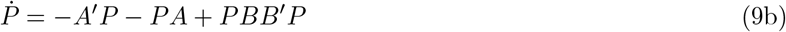

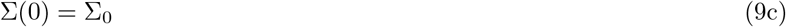

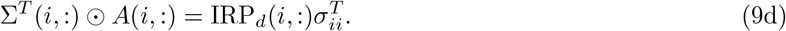

These results are summarized in Proposition 1, Supporting information C. Equation (9a) defines the evolution of the state covariance of the system in equation (6a) under the influence of the control input. The constraints in the evolution of the state covariance are defined by equations (9c) and (9d). It is important to highlight that equation (9b) shares similarities with a typical Linear Quadratic Regulator (LQR) problem, except the boundary constraint specified in (9d). The boundary value for *P* (*t*) in equation (9b) is unspecified and finding *P* (0), *P* (*T*) that satisfies the boundary constraints in (9d) is non-trivial. Also, the indefiniteness of *P* (*t*) places the context of our problem outside the standard LQR theory. Furthermore, the Riccati equations (9a) and (9b) are also coupled, not solely due to the boundary constraints, but also in terms of their dynamic behavior. Finding a closed-form solution to these coupled Riccati equations is challenging [57–62], and establishing both the existence and uniqueness of solutions for this system of equations is non-trivial. We summarize these arguments below.

*If there exists solutions, {P* (*t*), ∑(*t*)|0 ≤ *t* ≤ *T} that satisfy the coupled Riccati equations in (9a) and (9b) and the boundary conditions in equations (9c), (9d), then the optimal feedback gain in (8) solves Problem 7*.

In Theorem 2, Supporting information C, we show that there exist solutions *{P* (*t*), ∑(*t*)|0 ≤ *t* ≤ *T*} that satisfy the coupled Riccati equations in (9a) and (9b) provided 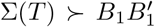. Further, we show that the state covariance can be steered between any two boundary constraints, thus satisfying the coupled boundary constraints in equations (9c) and (9d). Therefore, the set of control inputs *𝒰* that steers the IRP to the desired value is non-empty. To compute the optimal control inputs, we formulate the problem as a semidefinite program (SDP), which can be effectively solved using optimization tools like CVX [63] or YALMIP [64] as illustrated below.

**Numerical computation of optimal control:** Define the control input *u*(*t*) = −*K*(*t*)*ϕ*(*t*) so that

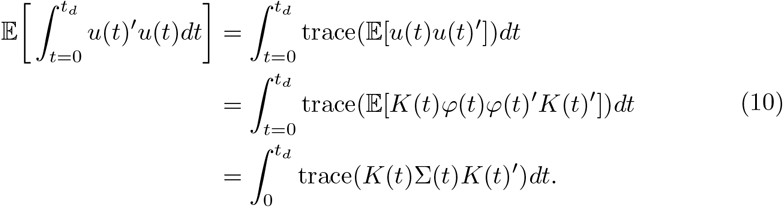

Define *U* (*t*) = −∑(*t*)*K*(*t*)′ such that the cost function in equation (10) can be written as

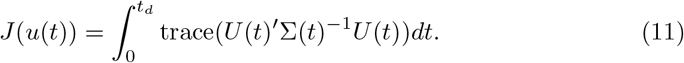

Thus, equation (11) is jointly convex in *U* (*t*) and ∑(*t*). Using *U* (*t*) = −∑(*t*)*K*(*t*)′, we can write equation (9a) as

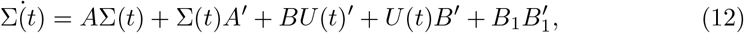

which is linear in both *U* (*t*) and ∑(*t*). The optimization problem in Problem (7) can thus be written as a semi-definite program as

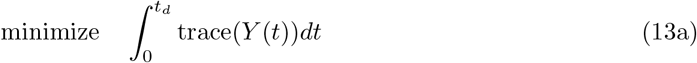

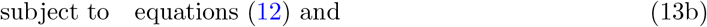

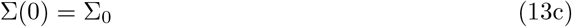

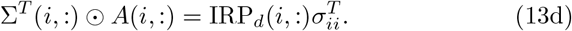

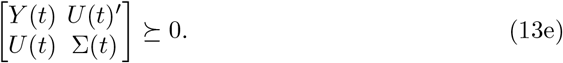

The problem can be solved by discretizing equation (12) and the feedback gain can be recovered as *K*(*t*) = −*U* (*t*)′∑(*t*)^−1^. Note that the SDP formulation in equation (13) can be employed to attain a desired pattern specifically for a subset of interacting nodes. For instance, if we require to achieve a desired IRP only from node *j* to node *i*, then the constraint in equation (13d) is replaced with the constraint 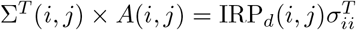.

#### 2.2.2 Minimum Energy Control to maintain stationary IRP

The objective here is to maintain a desired stationary IRP_*d*_ for the phase difference differential equation in (6a). Given that the pair (*A, B*) is controllable, the feedback control law is of the form *u*(*t*) = −*Kϕ*(*t*) (similar derivation as in Proposition 1, Supporting information C) and we are interested in the one that minimizes the input energy rate defined by *J*(*u*) = 𝔼 [*u*′*u*]. Similar to equation (7), given an admissible (Corollary 1.1 in Supporting information C) desired IRP_*d*_, we can formulate the minimum energy control problem as

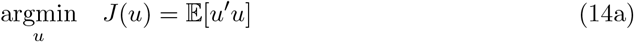

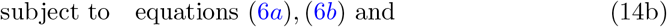

that equation (6a) admits IRP_*d*_ as the invariant IRP.

*The optimal control strategy u*^*^(*t*) *solving Problem (14) is of the form*

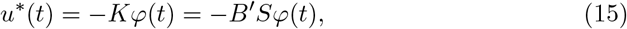

*where S is an n × n symmetric matrix such that* (*A* − *BB*′*S*) *is a Hurwitz matrix and satisfies both the following constraints:*

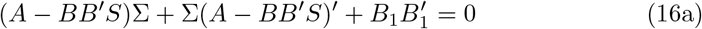

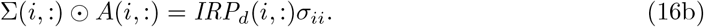

The proof is given in Theorem 3, Supporting information C. It is important to highlight that, in contrast to finite horizon control scenarios, the state covariance matrix ∑ remains invariant in this context. This invariant state covariance matrix serves as a key determinant of the desired stationary IRP for the oscillatory network via equation (16b).

**Numerical computation of optimal control:** Define the control input *u*(*t*) = −*Kϕ*(*t*) so that

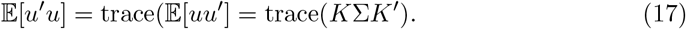

Define *M* = −∑*K*′ such that the cost function in equation (17) can be written as

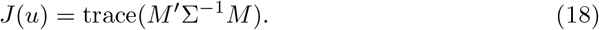

Thus, equation (18) is jointly convex in *M* and ∑. Using *M* = −∑*K*′, we can write equation (16a) as

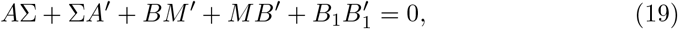

which is linear in both *M* and ∑. The optimization problem in Problem (9) can thus be written as a semi-definite program as

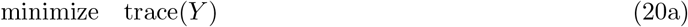

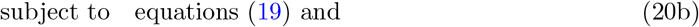

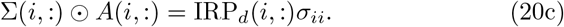

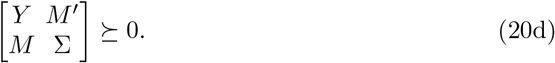

Similar to the finite horizon control, The problem can be solved by discretizing equation (19) feedback gain can be recovered as *K* = −*M* ′∑^−1^.

## 3 Applications

### 3.1 Gene Regulatory Networks

In this section, we consider a gene regulatory network consisting of two interconnected biochemical oscillators following the Goodwin-type dynamics (Supporting information D) as illustrated in Fig. 1a and 4a.

In a single oscillator, the gene (underlined rectangle) gets transcribed into mRNA (rectangle) that has concentration *x*_*i*_ within the cell. Subsequently, this mRNA is then translated into an enzyme (disk) with concentration *y*_*i*_. This enzyme helps in the production of a protein (triangle) with concentration *z*_*i*_. The concentration of the protein, in turn, suppresses the transcription of *x*_*i*_. This results in a nonlinear feedback loop that produces stochastic oscillatory dynamics as shown in Fig. 1b. The oscillatory dynamics are then reduced to the averaged phase stochastic dynamics of the form in equation (2). The fluctuating interactions between the two oscillators are then derived using the first-order approximation method (refer to Methods). The process reduces the nonlinear oscillatory network model into a simplified linear network dynamics model. The couplings between the two nodes are defined by the values of the coupling function at the stable phase locking states. The coupling function for gene regulatory model in Figs. 1a and 4a is shown in Fig. 2. The two oscillators exhibit two stable phase locking states and due to the anti-symmetric nature of the coupling function, the two stable states are separated by 2*π*. This leads to coupling values that are equal in magnitude but opposite in sign. To compute the information transfers, we assume that the initial covariance of the fluctuations is given by ∑(0) = **I**_2_, where **I**_2_ is an identity matrix of order 2. The evolution of the state covariance and the information transfers from time *t* = 0 to *t* = 120 are shown in Figures 4b-4d. The desired IRPs at *t* = {3, 50, 100} is given as |*T*_2→1_)(*t*)| = {0.001, 0.08, 0.0375}, |*T*_1→2_)(*t*)| = {0.001, 0.04, 0.0375} and the desired infinite horizon IRP is given as |*T*_1→2_(*t* = ∞)| = |*T*_2→1_(*t* = ∞)| = 0.0375, where |.| denotes the absolute values. We compute the optimal control inputs that drive the system towards the desired IRPs by solving equation (6). As we redirect the IRPs from a lower value to a higher value and then back to a lower value, we expect that fluctuations will progressively increase and then decrease gradually. The depicted phenomenon in Figure 4f illustrates the gradual increase in state covariance from *t* = 3 to *t* = 50, followed by a subsequent decrease until *t* = 100. In the infinite horizon scenario, when the IRP reaches a stable state, the state covariances also attain stability. Figures 4g and 4h show the evolution of the IRPs as they converge towards the predefined or desired IRPs during the time interval from *t* = 0 to *t* = 120.

**Fig 4.**
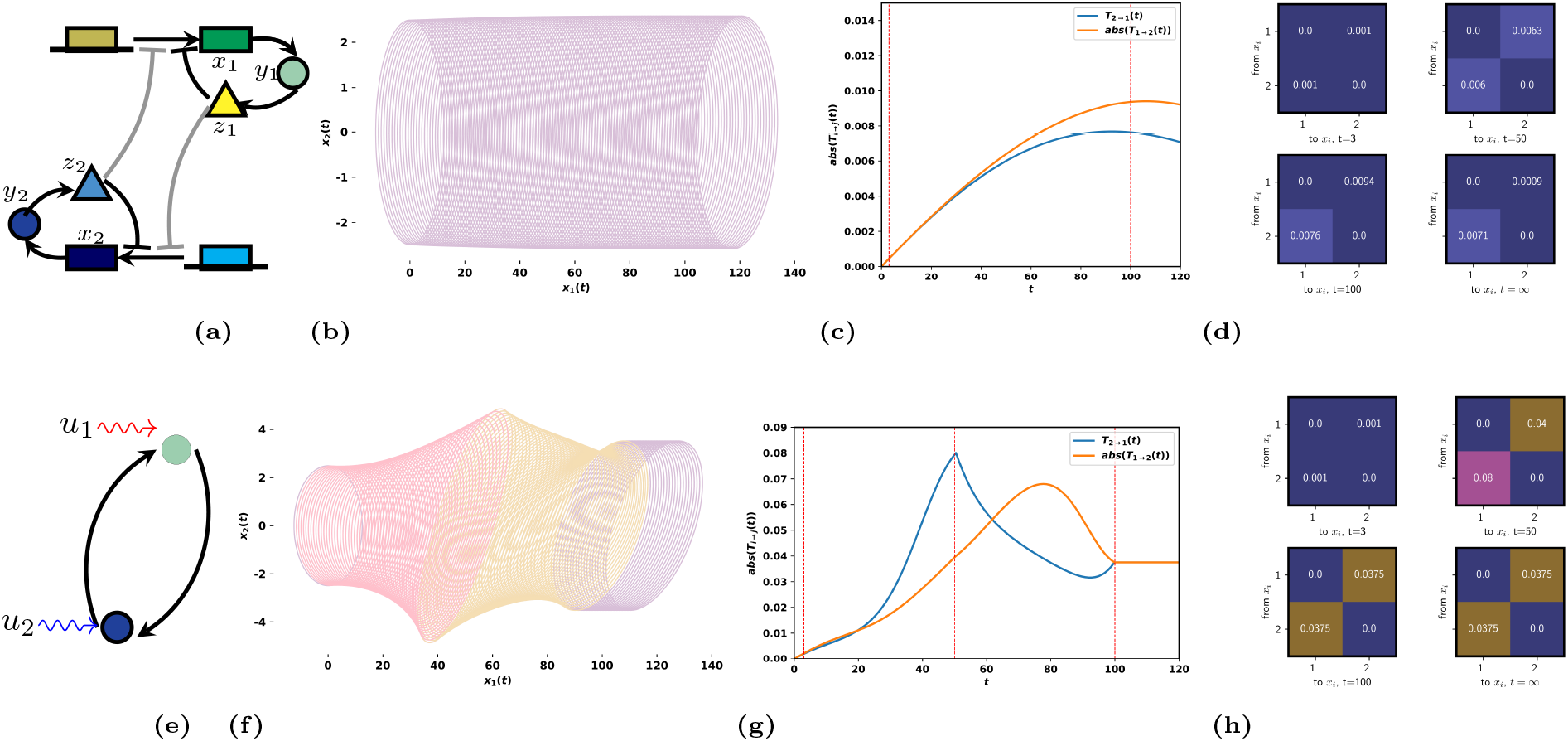
Finite and Infinite Horizon Control of IRPs: (a) Simple model of a gene-regulatory network of two coupled Goodwin oscillators (green and blue). Coupling strengths are gray coded (darker color indicates stronger coupling), sharp arrows indicate activating, and blunt arrows inhibiting influences. (b) The evolution of the state covariance from time *t* = 0 to *t* = 120 is illustrated by the time-varying ellipses. (c) Information flow curves between oscillators 1(*green*) to 2(*blue*) (d) Information routing patterns (IRPs) between the two oscillators at *t* = {3, 50, 100, 120}. (e) External input signals, *u*_1_(*t*) and *u*_2_(*t*) influence the linearized network dynamics model of fluctuating phase differences between the two oscillators. (f) Change in the evolution of the state distributions due to the influence of the external signals that reroute the IRPs to the desired IRPs as shown in Fig.(h) at *t* = {3, 50, 100} and at the infinite horizon. (g) Resulting transformation in the IRP curves at *t* = {3, 50, 100} as well as in the infinite horizon. (h) Attained target IRPs at target time instances.

### 3.2 Neurological Network

In this example, we study the information routing patterns among the excitatory populations of neurological networks. To model the dynamic interactions among the excitatory and inhibitory populations in a synaptically coupled neuronal network, we adopt the widely accepted Wilson-Cowan model of interacting oscillators (see Supporting information E). In neurological networks, a single neuron exhibits repeated firing when subjected to a constant current infection. Consequently, it is reasonable to regard a simulated neuron as a limit cycle, especially during short durations within a period of multiple spikes. We, thus, assume that each oscillator *i* has an asymptotically stable periodic solution characterized by a frequency denoted as *ω*_*i*_. The couplings among the neurons often consist of mild input currents affecting the membrane potential of the cells, suggesting the existence of weak couplings among the oscillators. We therefore, use averaging theory to derive equations that are solely dependent on the phase differences as in equation (2).

Figure 5 shows the phase reduction of a network of oscillatory dynamics consisting of 8 neurons. Supporting figure 1 shows the oscillatory dynamics of a pair of neurons along with the phase plane dynamics in the presence of stochastic noise. Examples of coupling functions and the anti-symmetric curves for two networks of 3 nodes are shown in Supporting figure 2. The stochastic component is approximated using linear approximations yielding a linear continuous stochastic model of the form in (28) (Methods and Supporting information B) and the corresponding reduced network model is shown in Figure 5d. We define the information transfer as the impact of one excitatory neuron on the excitation level of another neuron. The amount of information transfer depends on the degree of phase synchronization and the fluctuations around the stable phase synchronization state. In other words, information transfer is most effective when the pre-synaptic input of the sending neuron aligns with the post-synaptic neuron’s maximum excitability phase. We thus postulate that the increase in information transfer is a consequence of the increased fluctuations in the phase differences. A similar phenomenon can be noted in information transfers within gene regulatory networks, as illustrated in Figure 4f.

**Fig 5.**
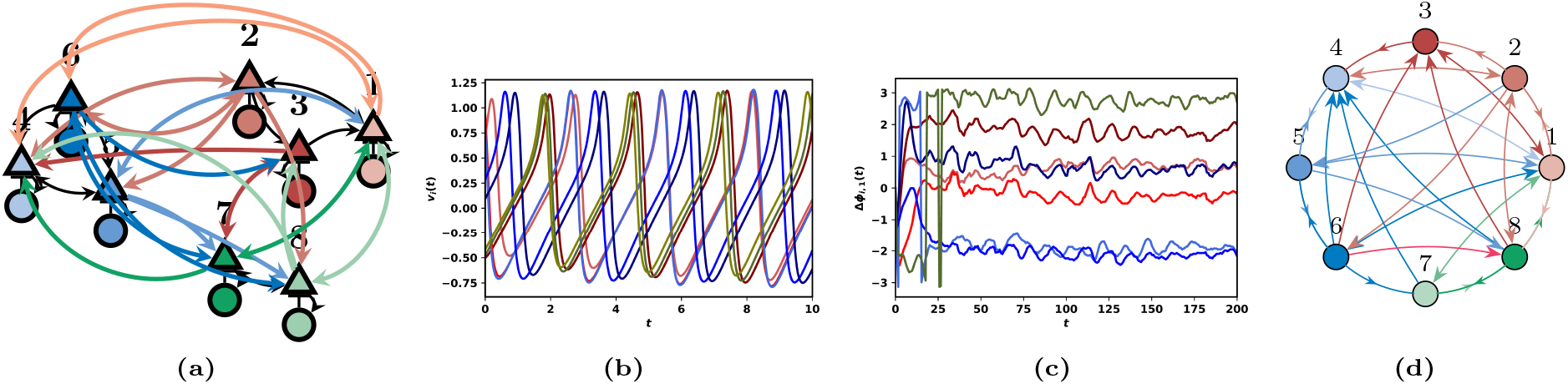
Phase reduction of a neuronal network consisting of excitatory (triangle) and inhibitory (disk) populations. (a) Excitatory and inhibitory network of 8 nodes. (b) Oscillatory behaviors of the excitatory neurons. (c) The oscillatory dynamics of the neurons exhibit fluctuating phase differences around a stable phase-locked state. (d) The reduced network that defines the fluctuations of the phase difference around the stable phase locking states.

Figure 6 illustrates the process involving the regulation of information flows across multiple excitatory neurons within a network model consisting of 8 neurons. We assume that the control nodes are nodes 1, 4, 6, and 8, as depicted in Figure 6h. The corresponding input matrix *B* is taken as *B* = [0.01 0 0 0.01 0 0.01 0 0.01]^*T*^. Additionally, a noise input matrix, denoted as *B*_1_, is assumed to be *B*_1_ = [0.01 0.01 0.01 0.01 0.01 0.01 0.01 0.01]′ and initial state covariance matrix is set as ∑(0) = 5**I**_8_. The heatmaps in Figures 6c and 6d represent the IRPs between all nodes in the network at *t* = 100, 200 respectively. The IRPs in Figures 6e and 6i show only the portions of the information transfer curves in the considered time interval. The minimum energy control inputs that direct the IRPs to the desired admissible patterns (as shown in Figure 6f) are computed by solving the SDP problem in Figure 6g. The constraint ∑(0) = ∑_0_ guarantees that the initial IRPs at *t* = 0 remain unaffected by the control actions. The IRP constraint in Figure 6g can be decomposed to indi-vidual IRPs as 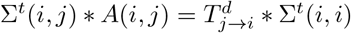, where 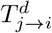 denotes the desired IRP from node *j* to node *i* at time *t*.

**Fig 6.**
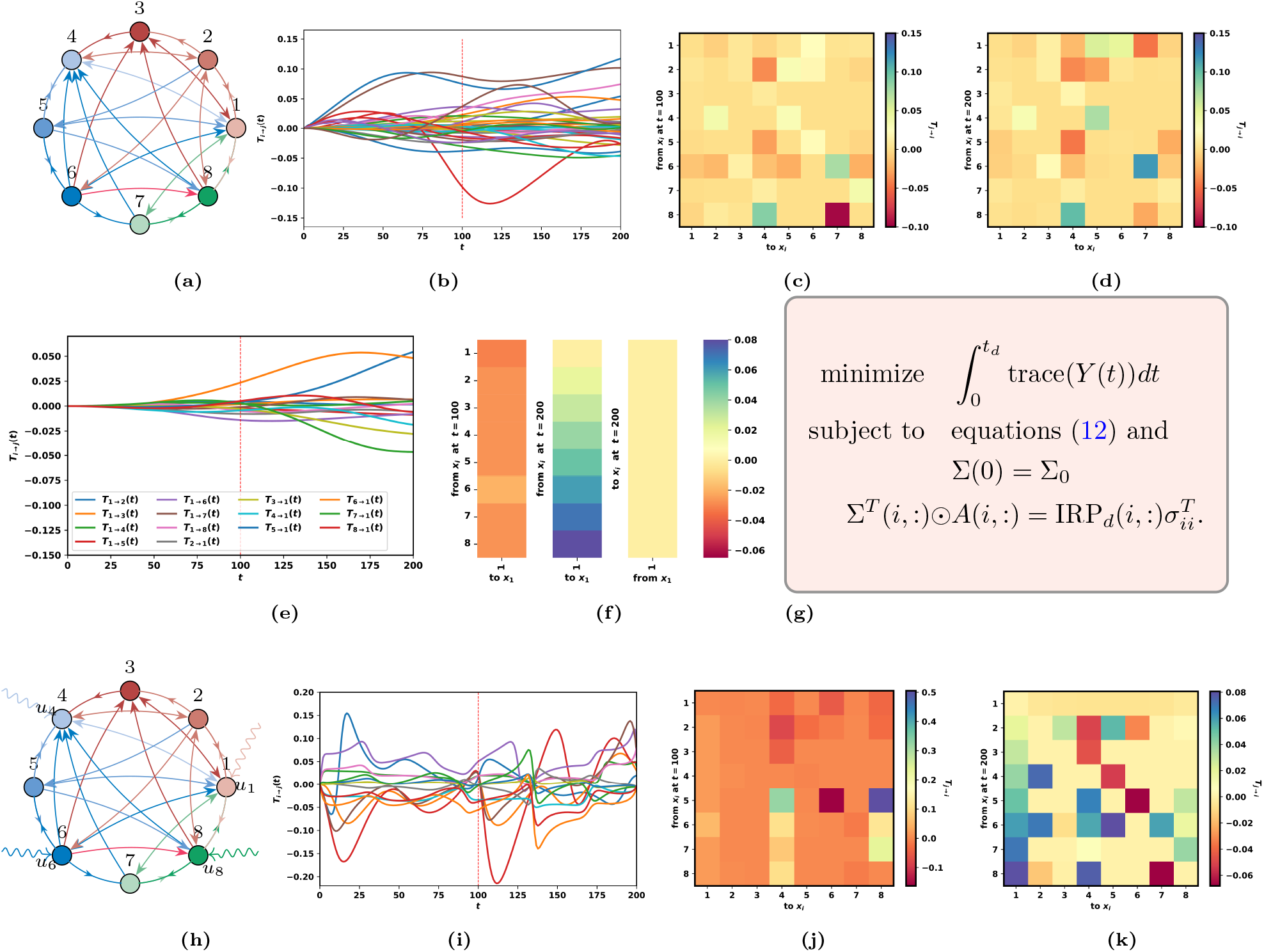
Control of IRPs of the coupled neuronal network in Fig. 5a. (a) The reduced phase difference network model as shown in Figure 5d. (b) The evolution of the IRPs from *t* = 0 to *t* = 200 (c) IRPs at *t* = 100 and (d) IRPs at *t* = 200 (e) The selected IRPs, *T*_*j*→*i*_ and the corresponding desired patterns are shown in (f) at *t* = 100 and *t* = 200. (g) The semi-definite programming to compute the control inputs that result in the desired IRPs. (h) The control inputs influencing the network dynamics (i) The evolutions of the controlled IRPs towards the desired patterns at *t* = 100 and *t* = 200. The achieved IRP_*d*_ can be observed at *t* = 100 and *t* = 200 in figures (j) and (k).

In the context of brain stimulation, nodes 1, 4, 6, and 8 are the stimulated nodes. The initial influences from the rest of the nodes to node 1 at *t* = 100 and *t* = 200 are shown in the first columns of the IRP matrices in Figures 6c and 6d. The desired influences from the rest of the nodes to node 1 under stimulation from nodes 1, 4, 6, and 8 at *t* = 100 are illustrated by the column matrix in Figure 6f. Figure 6j demonstrates the achieved IRP. In the second scenario, the goal is to gradually increase the influences from the rest of the nodes to node 1 from 0 to 0.08 under stimulation, while maintaining a negligible steady influence from node 1 to the rest of the nodes, as depicted in the second and third column matrices in Figure 6f. The resulting IRP under stimulation is shown in Figure 6k. The first column and the first row indicate that the desired stimulation pattern is achieved using optimal control inputs.

## 4 Discussion

The above results provide a fundamental basis for analyzing the information routing capabilities in complex dynamical networks with special focus in neurological and gene regulatory networks. The IRPs arise when the signals communicate due to fluctuations around stable reference states. Our study demonstrates how these patterns evolve in response to fluctuations, leading to varying routing patterns. We have discussed the joint effect of external perturbations, and noise on these IRPs.

Our results regarding these functional patterns are rooted in the fundamental concept of information transfer, which can be described as follows: *Information transfer from one random variable to another is quantified as the change in the entropy of the latter variable due to the influence of the former*. This foundational principle is indeed employed in deriving Kleeman’s information transfer as depicted in equation (23). Hence, this information-theoretic metric remains unassociated with specific algorithms or communication protocols, relying solely on the inherent dynamics of the network. For instance, consider Figure 1, where an external stimulation impacting the mRNA of one of the oscillators is represented through variations in the phase difference between the two oscillators. This is subsequently interpreted as a modification in the coupling function, ultimately leading to the emergence of two distinct routing patterns. Our theory is also based on a first-order approximation method, which also enables us to highlight the significance of the stable phase locking states within the underlying IRP. In network dynamical systems where higher-order interactions hold significance, the approximation can be systematically extended to incorporate these higher-order terms using the Perron-Frobenius operator [37].

The explicit dependence of the IRP on the underlying network dynamics motivates us to strategically manipulate the collective network dynamics to attain the intended IRP. For example, achieving the desired IRP might involve optimizing the network’s topology [24] or applying control theory techniques to determine the external signals that guide patterns toward the desired outcome. The principles in this study primarily focus on the application of control theory to address the latter challenge. To this end, we have introduced a problem formulation aimed at determining the minimum control energy required to guide information transfers to any desired value within both finite and infinite time horizons. Our analysis reveals that the problem can be addressed by identifying the control inputs needed to guide probability distributions toward predefined distributions. For example, it is shown in Figure 4 how the distribution in Figure 4b is steered as shown in Figure 4f to achieve the target IRP. We have established that the optimal inputs can be determined by solving coupled Riccati equations with coupled boundary constraints. However, it is important to note that no closed-form solutions exist, and we resort to nonlinear optimization tools such as CVX to find near-optimal control inputs. We demonstrated our theory with two biological networks (gene regulatory and neurological) where desired IRPs are required at desired instances.

## 5 Methods

### 5.1 Information Transfer Patterns in Network Dynamical Systems

To understand how the information transfer patterns change dynamically under the influence of dynamic nodal interactions in the network, consider a stochastic system where the dynamics are given by:

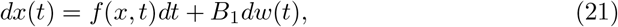

where *x*(*t*) ∈ R^*n*^ are the states of the system, *f* : ℝ^*n*^ *×* ℝ → ℝ^*n*^ describes the intrinsic network dynamics, *B*_1_ ∈ ℝ^*n×m*^ denotes the input noise matrix and *w*(*t*) ∈ ℝ^*m*^ is a white noise with mean zero and unit covariance. To quantify the instantaneous information flow from node *x*_*j*_ to node *x*_*i*_, we use an information-theoretic measure defined by ‘Information Transfer’ [37]. The formulation is based on the fundamental notion of information transfer. More precisely, for any two random variables, *x*_*i*_, *x*_*j*_ ∈ *x*(*t*), the transfer of information from *x*_*j*_ to *x*_*i*_ at time *t*, denoted as 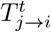 is defined as the variation in the instantaneous marginal entropy of *x*_*i*_ in the presence and the absence of the influence of *x*_*j*_. That is,

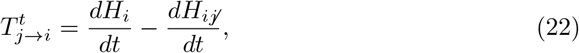

where 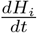 is the rate of change of marginal entropy of *x*_*i*_ from time *t* to *t* + Δ*t*, and 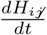 is the rate of change of marginal entropy of *x*_*i*_ with contributions from all other states except from *x*_*j*_. The information transfer from *x*_*j*_ to *x*_*i*_ is then derived from equation (22) as [37] (Supporting information A)

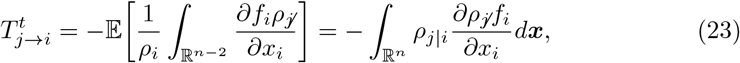

where 𝔼 denotes the expectation, *ρ*_*;j*_ denotes the joint distribution of (*x*_1_, *… x*_*j*−1_, *x*_*j*+1_, *… x*_*n*_) at time *t, ρ*_*i*_ denotes the marginal distribution of the state *x*_*i*_. and *w*(*t*) ∈ ℝ^*m*^ is a white noise with mean zero and unit covariance. The relationship between definition (23) and the widely used ‘Transfer Entropy’ is explained in Supporting information A. *T*_*j*→*i*_(*t*) is then computed for all edges originating from node *j* and reaching node *i*. This concept is analogous to functional connectivity and information routing patterns (IRPs) in biological networks. Hence, we refer to this as functional connectivity or IRPs.

### 5.2 Reduction to linear stochastic system

For a network stochastic dynamical system with *n* nodes, equation (2) can be written as

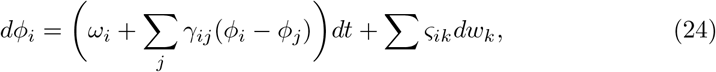

where *γ*_*ij*_(*ϕ*_*i*_ − *ϕ*_*j*_) are the coupling functions and external input are modeled as independent Wiener processes, *w*_*k*_. In the unperturbed system (*∑*_*ik*_ = 0), we assume that the phase dynamics described in equation (24) exhibit a stable phase-locked state with a consistent phase difference 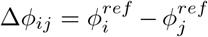 and a collective oscillation frequency Ω. This implies that for all *i* ∈ {1, *… n*},

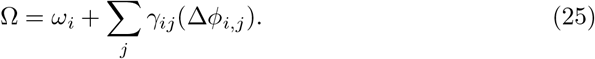

We decompose the phase dynamics into two components: a deterministic reference part denoted as 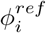 and a fluctuating part represented as 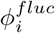. The solution to the deterministic dynamics is provided as follows:

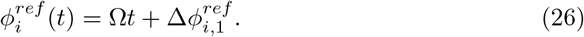

Introducing new coordinates, 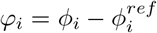, equation (24) can be written as

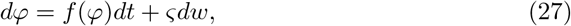

where 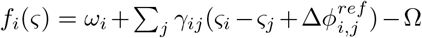. We assume that the noise levels, *∑*_*ik*_ are small, and using the small noise expansion, the first order approximation of equation (27) is given by a multivariate Ornstein-Uhlenbeck process

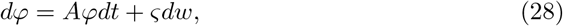

Where 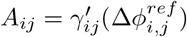. Thus, equation (28) is a linear stochastic continuous system, and the expression for *T*_*j*→*i*_(*t*) is given in equation (3).

## Acknowledgements

The work is partially supported by grants from Indo-US Science and Technology Forum (IUSSTF) IUSSTF/JC-110/2019.

## Author Contributions

SS: Conceptualization, Methodology, Writing – original draft. RP: Conceptualization, Resources, Writing – review & editing, Supervision, Funding acquisition. UV: Conceptualization, Visualization, Investigation. SL: Investigation, Writing – review & editing, Supervision.

## Supporting Information

- S1_file.pdf

## References

1. Volz, L.J., Sarfeld, A.-S., Diekhoff, S., Rehme, A.K., Pool, E.-M., Eickhoff, S.B., Fink, G.R., Grefkes, C.: Motor cortex excitability and connectivity in chronic stroke: a multimodal model of functional reorganization. Brain Structure and Function 220, 1093–1107 (2015)

2. Helfrich, C., Pierau, S.S., Freitag, C.M., Roeper, J., Ziemann, U., Bender, S.: Monitoring cortical excitability during repetitive transcranial magnetic stimulation in children with adhd: a single-blind, sham-controlled tms-eeg study. PloS one 7(11), 50073 (2012)

3. Tierney, T.S., Vasudeva, V.S., Weir, S., Hayes, M.T.: Neuromodulation for neurodegenerative conditions. Front. Biosci.(Elite. Ed) 5, 490–499 (2013)

4. DeGiorgio, C.M., Krahl, S.E.: Neurostimulation for drug-resistant epilepsy. Continuum: Lifelong Learning in Neurology 19(3), 743–755 (2013)

5. Rubio, B., Boes, A.D., Laganiere, S., Rotenberg, A., Jeurissen, D., Pascual-Leone, A.: Noninvasive brain stimulation in pediatric attention-deficit hyper-activity disorder (adhd) a review. Journal of child neurology 31(6), 784–796 (2016)

6. Hasan, A., Falkai, P., Wobrock, T.: Transcranial brain stimulation in schizophrenia: targeting cortical excitability, connectivity and plasticity. Current medicinal chemistry 20(3), 405–413 (2013)

7. Vanneste, S., Fregni, F., De Ridder, D.: Head-to-head comparison of transcranial random noise stimulation, transcranial ac stimulation, and transcranial dc stimulation for tinnitus. Frontiers in psychiatry 4, 158 (2013)

8. Lozano, A.M., Lipsman, N.: Probing and regulating dysfunctional circuits using deep brain stimulation. Neuron 77(3), 406–424 (2013)

9. Chaieb, L., Kovacs, G., Cziraki, C., Greenlee, M., Paulus, W., Antal, A.: Short-duration transcranial random noise stimulation induces blood oxygenation level dependent response attenuation in the human motor cortex. Experimental brain research 198, 439–444 (2009)

10. Steigerwald, F., Matthies, C., Volkmann, J.: Directional deep brain stimulation. Neurotherapeutics 16(1), 100–104 (2019)

11. Muldoon, S.F., Pasqualetti, F., Gu, S., Cieslak, M., Grafton, S.T., Vettel, J.M., Bassett, D.S.: Stimulation-based control of dynamic brain networks. PLOS Computational Biology 12(9), 1–23 (2016) 10.1371/journal.pcbi.1005076

12. Gu, S., Pasqualetti, F., Cieslak, M., Telesford, Q.K., Yu, A.B., Kahn, A.E., Medaglia, J.D., Vettel, J.M., Miller, M.B., Grafton, S.T., et al.: Controllability of structural brain networks. Nature communications 6(1), 8414 (2015)

13. Schüpbach, M., Chabardes, S., Matthies, C., Pollo, C., Steigerwald, F., Timmermann, L., Visser Vandewalle, V., Volkmann, J., Schuurman, P.R.: Directional leads for deep brain stimulation: Opportunities and challenges. Movement disorders 32(10), 1371–1375 (2017)

14. Pollo, C., Kaelin-Lang, A., Oertel, M.F., Stieglitz, L., Taub, E., Fuhr, P., Lozano, A.M., Raabe, A., Schüpbach, M.: Directional deep brain stimulation: an intraoperative double-blind pilot study. Brain 137(7), 2015–2026 (2014)

15. Tkačik, G., Callan Jr, C.G., Bialek, W.: Information flow and optimization in transcriptional regulation. Proceedings of the National Academy of Sciences 105(34), 12265–12270 (2008)

16. Tyson, J.J., Chen, K., Novak, B.: Network dynamics and cell physiology. Nature reviews Molecular cell biology 2(12), 908–916 (2001)

17. Kuramoto, Y., Kuramoto, Y.: Chemical Turbulence. Springer, ??? (1984)

18. Winfree, A.T.: The Geometry of Biological Time vol. 2. Springer, ??? (1980)

19. Strogatz, S.H.: Exploring complex networks. nature 410(6825), 268–276 (2001)

20. Pikovsky, A., Rosenblum, M., Kurths, J., Synchronization, A.: A universal concept in nonlinear sciences. Self 2, 3 (2001)

21. Acebrón, J.A., Bonilla, L.L., Vicente, C.J.P., Ritort, F., Spigler, R.: The kuramoto model: A simple paradigm for synchronization phenomena. Reviews of modern physics 77(1), 137 (2005)

22. Kirst, C., Timme, M., Battaglia, D.: Dynamic information routing in complex networks. Nature communications 7(1) (2016)

23. Menara, T., Baggio, G., Bassett, D., Pasqualetti, F.: Functional control of oscillator networks. Nature communications 13(1), 4721 (2022)

24. Singh, M.S., Pasumarthy, R., Vaidya, U., Leonhardt, S.: On quantification and maximization of information transfer in network dynamical systems. Scientific Reports 13(1), 5588 (2023)

25. Battaglia, D., Witt, A., Wolf, F., Geisel, T.: Dynamic effective connectivity of inter-areal brain circuits. PLoS computational biology 8(3), 1002438 (2012)

26. Bennett, S.H., Kirby, A.J., Finnerty, G.T.: Rewiring the connectome: evidence and effects. Neuroscience & Biobehavioral Reviews 88, 51–62 (2018)

27. Gu, S., Betzel, R.F., Mattar, M.G., Cieslak, M., Delio, P.R., Grafton, S.T., Pasqualetti, F., Bassett, D.S.: Optimal trajectories of brain state transitions. Neuroimage 148, 305–317 (2017)

28. Raichle, M.E.: The brain’s default mode network. Annual review of neuroscience 38, 433–447 (2015)

29. Baggio, G., Bassett, D.S., Pasqualetti, F.: Data-driven control of complex networks. Nature communications 12(1), 1429 (2021)

30. Liu, Y.-Y., Slotine, J.-J., Barabási, A.-L.: Controllability of complex networks. nature 473(7346), 167–173 (2011)

31. Campbell, C., Ruths, J., Ruths, D., Shea, K., Albert, R.: Topological constraints on network control profiles. Scientific reports 5(1), 18693 (2015)

32. Gutiérrez, R., Sendina-Nadal, I., Zanin, M., Papo, D., Boccaletti, S.: Targeting the dynamics of complex networks. Scientific reports 2(1), 396 (2012)

33. Yuan, Z., Zhao, C., Di, Z., Wang, W.-X., Lai, Y.-C.: Exact controllability of complex networks. Nature communications 4(1), 2447 (2013)

34. Kim, J.Z., Soffer, J.M., Kahn, A.E., Vettel, J.M., Pasqualetti, F., Bassett, D.S.: Role of graph architecture in controlling dynamical networks with applications to neural systems. Nature physics 14(1), 91–98 (2018)

35. Pósfai, M., Liu, Y.-Y., Slotine, J.-J., Barabási, A.-L.: Effect of correlations on network controllability. Scientific reports 3(1), 1067 (2013)

36. San Liang, X., Kleeman, R.: Information transfer between dynamical system components. Physical review letters 95(24), 244101 (2005)

37. San Liang, X., Kleeman, R.: A rigorous formalism of information transfer between dynamical system components. i. discrete mapping. Physica D: Nonlinear Phenomena 231(1), 1–9 (2007)

38. Asafo-Adjei, E., Owusu Junior, P., Adam, A.M.: Information flow between global equities and cryptocurrencies: a vmd-based entropy evaluating shocks from covid-19 pandemic. Complexity 2021, 1–25 (2021)

39. Asafo-Adjei, E., Frimpong, S., Owusu Junior, P., Adam, A.M., Boateng, E., Ofori Abosompim, R.: Multi-frequency information flows between global commodities and uncertainties: evidence from covid-19 pandemic. Complexity 2022, 1–32 (2022)

40. Hagan, D.F.T., Wang, G., San Liang, X., Dolman, H.A.: A time-varying causality formalism based on the liang–kleeman information flow for analyzing directed interactions in nonstationary climate systems. Journal of Climate 32(21), 7521–7537 (2019)

41. Ay, N., Polani, D.: Information flows in causal networks. Advances in complex systems 11(01), 17–41 (2008)

42. Sinha, S., Vaidya, U.: On data-driven computation of information transfer for causal inference in discrete-time dynamical systems. Journal of Nonlinear Science 30(4), 1651–1676 (2020)

43. Ermentrout, G.B., Kopell, N.: Multiple pulse interactions and averaging in systems of coupled neural oscillators. Journal of Mathematical Biology 29(3), 195–217 (1991)

44. Pietras, B., Daffertshofer, A.: Network dynamics of coupled oscillators and phase reduction techniques. Physics Reports 819, 1–105 (2019)

45. Zhang, F., Leonard, N.E.: Coordinated patterns of unit speed particles on a closed curve. Systems & control letters 56(6), 397–407 (2007)

46. Buck, J., Buck, E.: Biology of synchronous flashing of fireflies. Nature Publishing Group UK London (1966)

47. Dörfler, F., Chertkov, M., Bullo, F.: Synchronization in complex oscillator networks and smart grids. Proceedings of the National Academy of Sciences 110(6), 2005–2010 (2013)

48. Trenberth, K.E.: Spatial and temporal variations of the southern oscillation. Quarterly Journal of the Royal Meteorological Society 102(433), 639–653 (1976)

49. Cabral, J., Hugues, E., Sporns, O., Deco, G.: Role of local network oscillations in resting-state functional connectivity. Neuroimage 57(1), 130–139 (2011)

50. Fell, J., Axmacher, N.: The role of phase synchronization in memory processes. Nature reviews neuroscience 12(2), 105–118 (2011)

51. Váša, F., Shanahan, M., Hellyer, P.J., Scott, G., Cabral, J., Leech, R.: Effects of lesions on synchrony and metastability in cortical networks. Neuroimage 118, 456–467 (2015)

52. Menara, T., Lisi, G., Pasqualetti, F., Cortese, A.: Brain network dynamics fingerprints are resilient to data heterogeneity. Journal of Neural Engineering 18(2), 026004 (2021)

53. González-García, C., Flounders, M.W., Chang, R., Baria, A.T., He, B.J.: Content-specific activity in frontoparietal and default-mode networks during prior-guided visual perception. elife 7, 36068 (2018)

54. Feillet, C., Krusche, P., Tamanini, F., Janssens, R.C., Downey, M.J., Martin, P., Teboul, M., Saito, S., Lévi, F.A., Bretschneider, T., et al.: Phase locking and multiple oscillating attractors for the coupled mammalian clock and cell cycle. Proceedings of the National Academy of Sciences 111(27), 9828–9833 (2014)

55. Purvis, J.E., Lahav, G.: Encoding and decoding cellular information through signaling dynamics. Cell 152(5), 945–956 (2013)

56. Katsuhiko, O.: Modern Control Engineering. Editorial Félix Varela, ??? (2009)

57. Chen, Y., Georgiou, T.T., Pavon, M.: Optimal steering of a linear stochastic system to a final probability distribution, part ii. IEEE Transactions on Automatic Control 61(5), 1170–1180 (2015)

58. Chen, Y., Georgiou, T.T., Pavon, M.: Optimal steering of a linear stochastic system to a final probability distribution, part i. IEEE Transactions on Automatic Control 61(5), 1158–1169 (2015)

59. Wakolbinger, A., et al.: Schrödinger bridges from 1931 to 1991. In: Proc. of the 4th Latin American Congress in Probability and Mathematical Statistics, Mexico City, pp. 61–79 (1990)

60. Beurling, A.: An automorphism of product measures. Annals of Mathematics, 189–200 (1960)

61. Jamison, B.: Reciprocal processes: The stationary gaussian case. The Annals of Mathematical Statistics 41(5), 1624–1630 (1970)

62. Georgiou, T.T., Pavon, M.: Positive contraction mappings for classical and quantum schrödinger systems. Journal of Mathematical Physics 56(3) (2015)

63. Grant, M., Boyd, S.: CVX: Matlab software for disciplined convex programming, version 2.1 (2014)

64. Lofberg, J.: Yalmip: A toolbox for modeling and optimization in matlab. In: 2004 IEEE International Conference on Robotics and Automation (IEEE Cat. No. 04CH37508), pp. 284–289 (2004). IEEE

